# Combined effect of matrix quality and spatial heterogeneity on biodiversity decline

**DOI:** 10.1101/324624

**Authors:** Irene Ramos, Cecilia González González, Ana L. Urrutia, Emilio Mora Van Cauwelaert, Mariana Benítez

## Abstract

The land-sparing/land-sharing debate remains an oversimplified framework to evaluate landscape management strategies that aim to reconcile food production and biodiversity conservation. Still, biodiversity-yield curves, on which the framework has relied, provide valuable qualitative information on biodiversity’s sensitivity to agricultural practices, and much research has studied this relationship. But the potential effect of landscape configuration on biodiversity’s response to intensification has rarely been considered; besides, studies have often taken yield as an indicator of agricultural management and have generalized conclusions from studying particular taxonomic groups. In this work we adapt a metacommunity model to analyze factors that shape biodiversity’s response to agricultural intensification while addressing some of the simplifications of the land-sparing/land-sharing dichotomy. In particular, we study species richness decline in landscapes encompassing a combined gradient in matrix quality and configurational heterogeneity of habitat patches, and considering community dynamics. We found that species richness along an intensification gradient shifts from following a robust response to presenting an abrupt decline, as landscape heterogeneity increases, as stricter survival thresholds are applied to species or as habitat area is reduced. Our work highlights the interdependent effects of heterogeneity, habitat availability and matrix quality on biodiversity and contributes to a nuanced understanding of ecological and landscape factors that enable a robust response of biodiversity in the face of spatiotemporal perturbations.

## INTRODUCTION

In the context of global food insecurity and biodiversity loss, it is critical to develop landscape management strategies that address these interrelated problems sustainably. The land-sparing/land-sharing debate has brought attention to this issue, framing it as a dichotomy between management strategies. On one hand, land-sparing proposes to limit food production to a small surface, where high-intensity agriculture is practiced to maximize yield, while other zones are reserved for biodiversity conservation. Therefore, landscapes associated with this strategy tend to have low spatial heterogeneity and a low quality matrix, that is, habitat patches embedded in so-called intensified plots where local biodiversity is unlikely to thrive (Fischer et al. 2008, Fischer et al. 2014, Kremen 2015). On the other hand, land-sharing proposes to use wildlife-friendly agriculture at the expense of a bigger production surface and to integrate agricultural areas with areas for conservation throughout the landscape; as a result, landscapes tend to maintain a high-quality matrix and high heterogeneity (Fischer et al. 2008, Fischer et al. 2014, Kremen 2015). Land-sparing and land-sharing strategies can thus be thought as opposites in a combined gradient of two variables: matrix quality and spatial heterogeneity. However, most empirical and theoretical efforts to compare such strategies focus on either of these variables, without explicitly considering their combined effect on biodiversity (Villard and Metzger 2014, Butsic and Kuemmerle 2015, Chaplin-Kramer et al. 2015, Liao et al. 2016). Indeed, in a previous work, we focused only on the effect of matrix intensification on biodiversity conservation (González González et al., 2016).

The land-sparing/land-sharing framework has largely relied on biodiversity-yield curves to contrast management strategies, as their shapes provide insight on the impact of agricultural practices on species persistence (Perfecto et al. 2009, Phalan et al. 2011, Butsic and Kuemmerle 2015, González González et al. 2016). If biodiversity declines drastically as management intensity increases, that is, following a concave up curve, then it is argued that a land-sparing approach would minimize biodiversity loss by minimizing the managed area. If, instead, biodiversity declines following a concave down curve, suggesting a robust response to low intensity agriculture, then a land-sharing approach is preferred. From a theoretical perspective, this qualitative information is valuable in evaluating biodiversity’s sensitivity to agricultural intensification.

However, it is recognized that the land-sparing/land-sharing framework oversimplifies the tradeoffs involved in achieving food security and sovereignty (Chapell and LaValle 2011, Fischer et al. 2011, Kremen 2015, Bennett 2017). The debate has centered on the tradeoff between yield and biodiversity conservation, often taking agricultural production as an indicator for agricultural management. For example, a common assumption is that less intense agricultural practices necessarily produce lower yields, in spite of evidence that they could satisfy global food demand (Chappell and LaValle 2011, Tscharntke et al. 2012). Instead, it has been proposed that determining biodiversity’s responses with respect to agricultural management is more pertinent in evaluating conservation strategies and in discussing food sovereignty beyond productivity (Perfecto et al. 2009, Kremen 2015). We focus on this relationship, considering the intensification of agricultural management as the “transition from ecosystems with high planned biodiversity and a more traditional management style to low planned biodiversity and an industrial management style, such as the use of agrochemicals” (Perfecto et al. 2009:19).

Another limitation of this framework is that the spatial scale and configuration at which habitat patches should be preserved is not specified. While land-sparing strategies intend to preserve extensive and continuous habitat patches, as opposed to the small and dispersed fragments often associated with land-sharing, the heterogeneity that results from these spatial arrangements has not been defined consistently (Fischer et al. 2014, Kremen 2015). The lack of an explicit distinction between configurational heterogeneity (spatial arrangement of cover types) and compositional heterogeneity (the number and proportions of cover types) contributes to the confusion (Fahrig et al. 2011); so, when land-sharing is said to promote higher heterogeneity than land-sparing, it is not always clear which aspect of heterogeneity it refers to. Still, configurational heterogeneity has been the most neglected, to the extent that studying biodiversity’s response to an intensification gradient (i.e. a reduction in the number of cover types), as exemplified by typical biodiversity-yield curves, accounts for the compositional aspect of heterogeneity. We consider the effect on biodiversity of configurational heterogeneity of habitat patches together with an intensification gradient (Figure 1); this component of heterogeneity has mainly been studied outside the land-sparing/land-sharing framework, with an emphasis on habitat loss over fragmentation or matrix quality (Fahrig 2003; Liao et al. 2016).

Finally, the inconsistency of indicators used in assessing biodiversity’s response to intensification is another limitation that concerns us. Comparisons of land-sparing/land-sharing strategies in empirical studies have mostly focused on particular taxonomic groups or populations instead of ecological communities (Prevedello and Vieira 2010, Phalan et al. 2011). Studies have also tended to measure a variety of ecological variables that may not be a direct indicator of biodiversity persistence, for example, by focusing only on abundance, or that may not be comparable among studies. These difficulties have hindered progress on the debate.

In a previous work, González González and collaborators (2016) developed a spatially explicit metacommunity model to study the impact of matrix quality on biodiversity. They found that richness declines following a concave down curve as matrix quality decreases, suggesting a robust response of biodiversity to agricultural intensification; however, the study did not consider the potential effect of different landscape configurations, among other factors that may shape biodiversity-loss curves. Here we adapt this model to study the shape of richness decline in a combined gradient of matrix quality and spatial heterogeneity, and we test the effect of habitat proportion and populations´ survival thresholds on the curves’ shapes. We found that as particular stresses on biodiversity increase (fragmentation, habitat loss and population vulnerability), richness decline along an intensification gradient shifts from a concave down to a concave up shape.

## METHODS

We adapt a metacommunity model (González González et al. 2016) to study the combined effect of matrix quality and heterogeneity on biodiversity, as well as the impact of habitat proportion and populations’ survival thresholds. This model simulates the spatial distribution of a hypothetical ecological community in an agricultural landscape by coupling a local community network dynamic with a migration dynamic. We present an overview of the model and the adaptations we implemented.

### Communities

Using the niche model of food webs (Williams and Martinez 2000), we generated 100 networks of 10 trophic species. From these we built interaction matrices by assigning random weights to edges and set up a Lotka-Volterra system with which we model species interactions. For each community, the system has the form

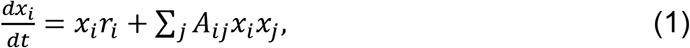

where *x*_*i*_ is the abundance of species *i*, *r*_*i*_ is the intrinsic growth rate of species *i*, and *A*_*ij*_ is the interaction matrix derived from the niche model. We parameterized the system choosing random initial abundances and growth rates. Although these communities are a subset from those used in our previous work (González González et al. 2016), we adjusted parameters to achieve higher average coexistence; this allows us to measure a wider range in species richness.

### Landscape

We simulate landscapes with a lattice of 10 by 10 cells with periodic boundary. Each cell represents one patch of either primary vegetation (or *habitat*), *high-quality agriculture* or *low-quality agriculture*, and may be occupied by a community, that is, interacting populations of multiple species. We define the *quality of the matrix* as the proportion of high-quality agriculture patches; thus replacing high-quality agriculture patches for low-quality ones models a decrease in matrix quality or *intensification*.

In addition, we define *landscape heterogeneity* as the total edge between habitat patches and high-quality or low-quality agriculture patches (See vertical axis on Figure 1). We map this metric to five qualitative heterogeneity levels to ensure comparability between landscapes with different habitat area, so that for a given amount of habitat, we consider the lowest possible heterogeneity (when all habitat patches are clustered and contiguous) as level 0 and the highest possible heterogeneity (when habitat patches are apart and distributed throughout the landscape) as level 4.

In our experiments we simulate biodiversity distribution in landscapes along ten degrees of intensification and five levels of heterogeneity (Figure 1). To generate these landscapes we specified the location of habitat patches, verifying that they were sparsely distributed, and chose random positions for the other two types of patches.

**Figure 1.**
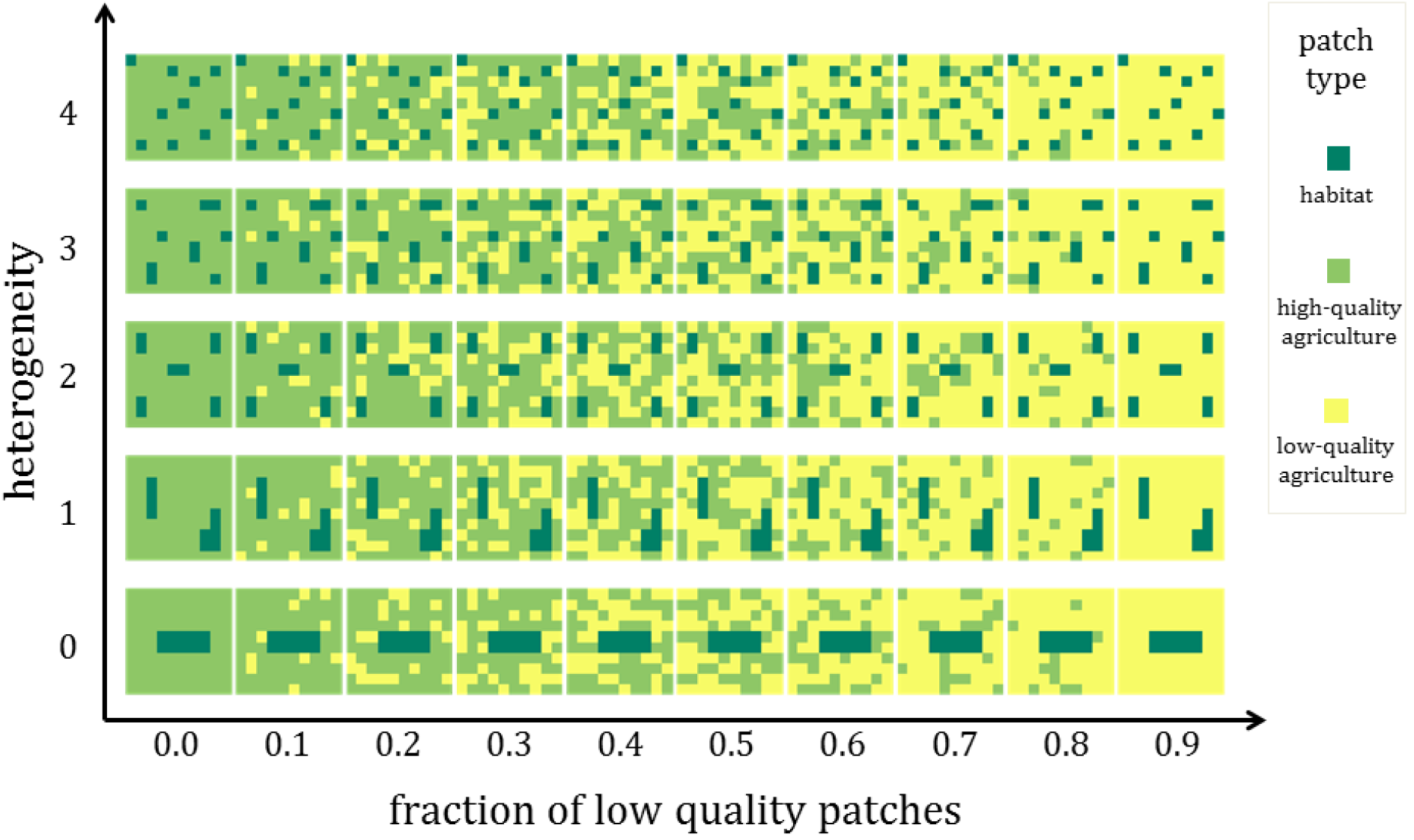
Combined gradient in matrix quality and heterogeneity. Hypothetical landscapes with 10% habitat area, in a combined gradient of matrix quality (as measured by the fraction of low-quality patches in the x-axis) and spatial heterogeneity (as measured by total habitat edge in the y-axis).

### Simulation

The model takes as entries a community (characterized by its interaction matrix, species’ intrinsic growth rates and initial abundances) and a landscape. On initialization, only habitat patches are occupied by a community, with specified initial abundances. The model iterates between community dynamics and migration until arriving to a steady state, for which we found 100 time steps are sufficient (Figure 2).

During each time step: i) Communities in habitat patches interact as described by the previous Lotka-Volterra system (Equation 1). Then, ii) a proportion of each population, in all patches, migrates to the eight neighboring patches depending on the type of patch: 30% in habitat patches and 100% in high-quality or low-quality agriculture patches; next, in all patches, a proportion of incoming populations dies as a cost of migration, again, depending on the type of patch: 0% in habitat patches, 30% in high-quality agriculture and 85% in low-quality agriculture. Coupling community and migration dynamics requires a given number *n* of migration substeps for every community dynamics substep, as these processes occur at different time scales; in our simulations *n* = 5.

The model’s output is the spatial distribution of species across the landscape in steady state, to which we can apply richness or abundance measures. Here we focus on species richness, defined as the number of species whose final abundance in the landscape is larger than an arbitrary *survival threshold*. We interpret this parameter as the minimum viable population, which is the smallest isolated population with a chance of surviving despite potential demographic, environmental or genetic stochasticity (Shaffer 1981).

**Figure 2.**
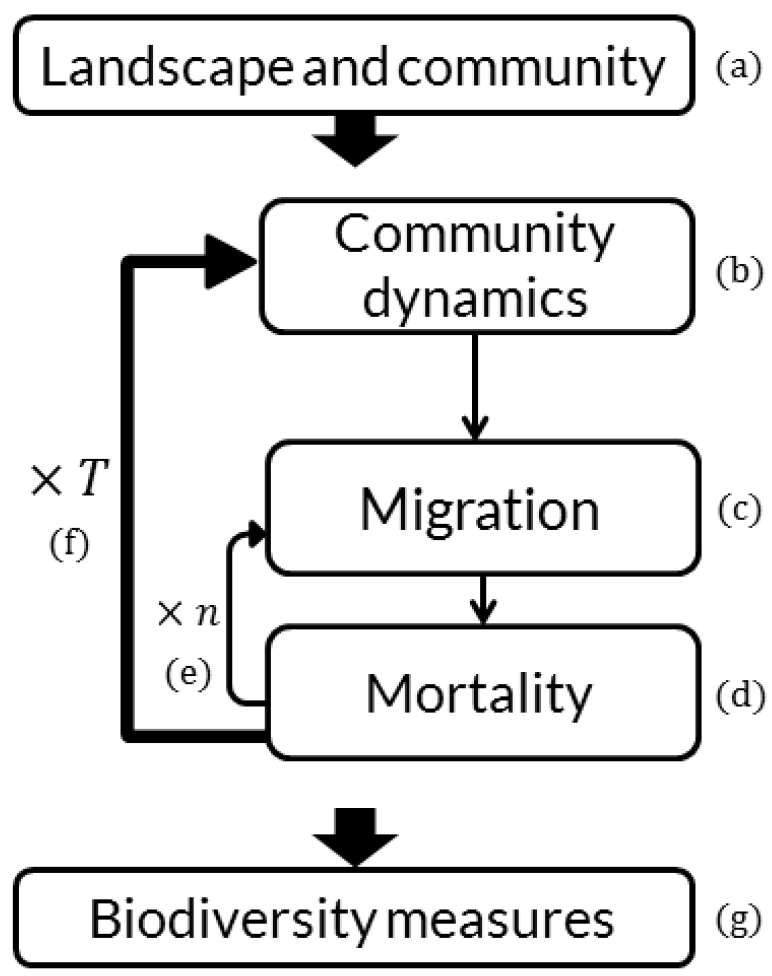
Model diagram. Diagram of the implemented metacommunity model (adapted from González González et al. 2016). The model takes as entries a landscape and an ecological community (a); it iterates between a community dynamic (b) modeled as a generalized Lotka-Volterra system and a migration dynamic (c) that includes a mortality substep (d) Coupling both dynamics requires *n* substeps of migration and mortality for every community dynamic substep (e); we use *n* = 5. The simulation lasts *T* time steps until arriving to a steady state (f); we use *T* = 1 0 0. Richness is calculated from the resulting spatial distribution of species across the landscape (g).

### Experiments and statistical analyses

We simulate biodiversity distribution in landscapes with 10% and 30% of habitat patches, and in each case we consider five heterogeneity levels and ten degrees of intensification, for a total of 50 landscapes per habitat area (Figure 1). We run the model in every landscape with 100 hypothetical communities and measure the resulting species richness. Given that mean abundance varies by orders of magnitude among communities, we restrict the analyses to communities that can be compared applying the same survival threshold; specifically, we select 79 communities that, at an intermediate heterogeneity level in 10%-habitat landscapes, show a change in richness along the intensification gradient under a *survival threshold* = 30. At every combination of habitat area, heterogeneity and intensification, we calculate mean species richness and standard deviation, then we study the shape of mean richness decline as matrix quality decreases in each heterogeneity level and the differences in richness between heterogeneity levels.

In particular, at each heterogeneity level we fit first and second degree polynomials to mean species richness along the intensification gradient using the least-squares method and choose the best fit as the one with higher adjusted *R*^2^. In the case of second degree polynomials, we characterize the shape of the curves by their quadratic term coefficient: a negative coefficient indicates that biodiversity decreases as a concave down curve, while a positive coefficient indicates it decreases as a concave up curve (González González et al. 2016). We identify differences in richness between heterogeneity levels by computing Kruskal-Wallis tests at every degree of intensification.

### Implementation

The model, simulations, data analyses and figure creation were implemented in the Python programming language, using the libraries NumPy, SciPy, Matplotlib and Seaborn. The scripts are available in a version-control repository at [https://github.com/laparcela/MatrizAgroecologica].

## RESULTS

Richness decline along an intensification gradient shifts from a concave down to a concave up shape as stresses on biodiversity increase, either by augmenting landscape heterogeneity (specifically, configurational heterogeneity measured as total habitat edge), applying stricter survival thresholds to species or reducing habitat area.

### Effect of heterogeneity on the shape of richness decline curves

For each heterogeneity level in 10%-habitat landscapes (Figure 3, Supplementary Figure 1), we first determine whether richness declines linearly or nonlinearly. Here we use *survival threshold* = 30. In landscapes with the lowest heterogeneity (level 0), richness declines as a concave down curve as matrix quality decreases. Next, in heterogeneity level 1, richness declines linearly with a relatively mild slope. Last, in high heterogeneity landscapes (levels 2, 3 and 4), richness declines as a concave up curve. This suggests that richness decline along an intensification gradient shifts from a concave down to a concave up shape as landscape heterogeneity increases.

**Figure 3.**
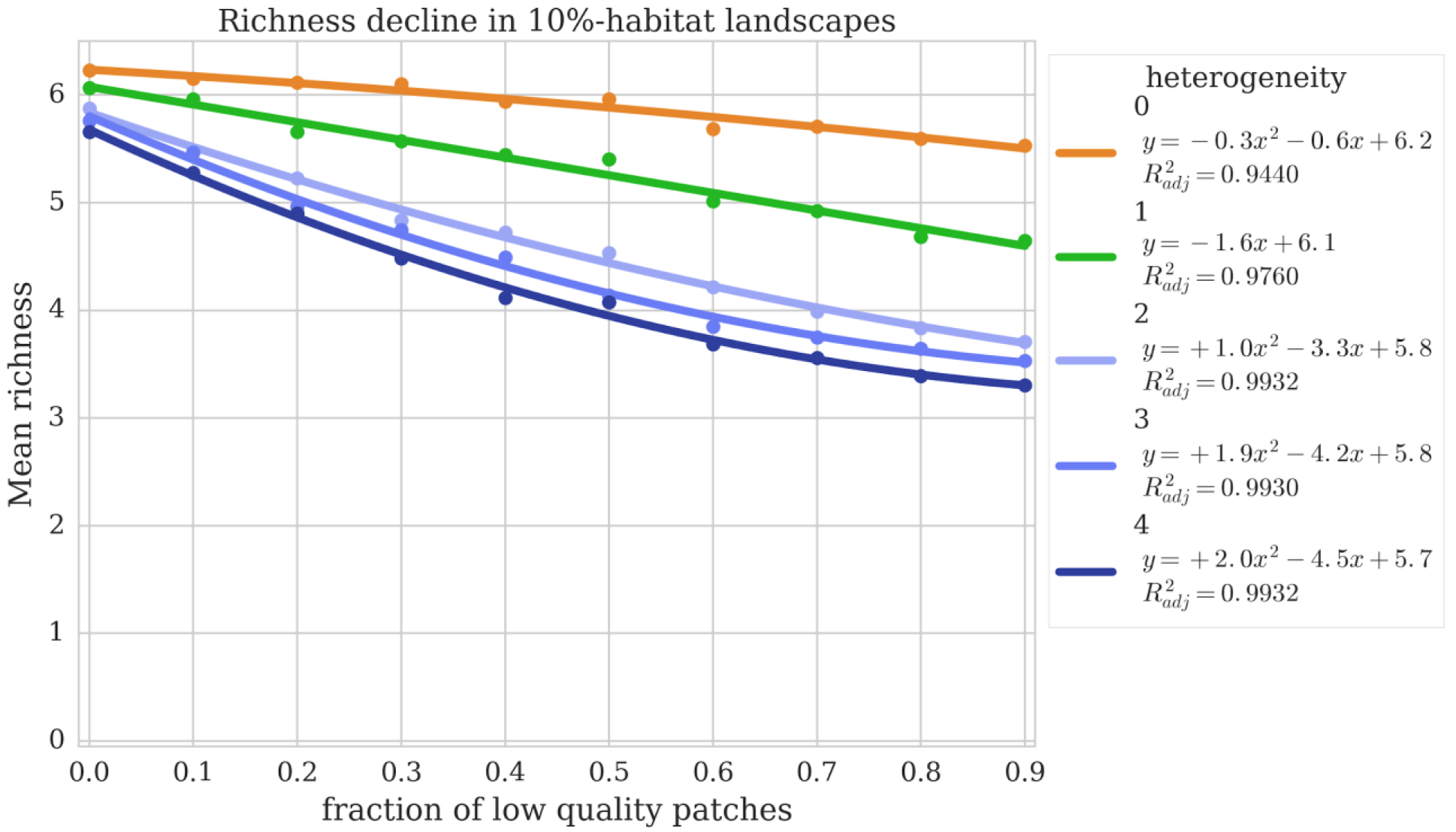
Richness decline in 10%-habitat landscapes. Richness decline along an intensification gradient (x-axis) in 10%-habitat landscapes (*survival threshold* = 30). In low heterogeneity landscapes (level 0), richness declines as a concave down curve, suggesting a robust response of biodiversity to intensification; in contrast, richness follows a concave up curve in high heterogeneity landscapes (levels 4, 3, and 2). Each data point is the average richness of 79 communities; see Supplementary Figure 1 for full distributions.

### Differences in richness between heterogeneity levels

As matrix quality decreases, richness remains higher in low heterogeneity landscapes than in high heterogeneity landscapes, but at low intensification heterogeneity does not have a considerable impact on richness. High heterogeneity landscapes have similar richness at each degree of intensification (blue curves in Figure 3). A Kruskal-Wallis test computed at every degree of intensification showed no difference between the distribution of species richness in landscapes with heterogeneity levels 4, 3 and 2 (p>0.05 in all cases).

However, along the intensification gradient landscapes at the two extremes of heterogeneity have contrasting richness. A second Kruskal-Wallis test computed at every degree of intensification showed a significant difference in the distribution of richness between landscapes with the highest and lowest heterogeneity, that is, level 4 compared with level 0 (p<0.05 in all cases), except at 0% of intensification.

Finally, we determined where along the gradient there are three statistically distinct responses to heterogeneity levels (Figure 4a). We compared, via Kruskal-Wallis tests, richness distribution between (i) heterogeneity levels 4 and 1, and (ii) heterogeneity levels 1 and 0, at every degree of intensification. The tests showed significant differences in both comparisons (p<0.05) from 60% to 90% of intensification, implying that within that interval, richness distributions in landscapes with heterogeneity levels 4, 1 and 0 (blue, green and orange curves, respectively) are distinguishable from each other.

**Figure 4.**
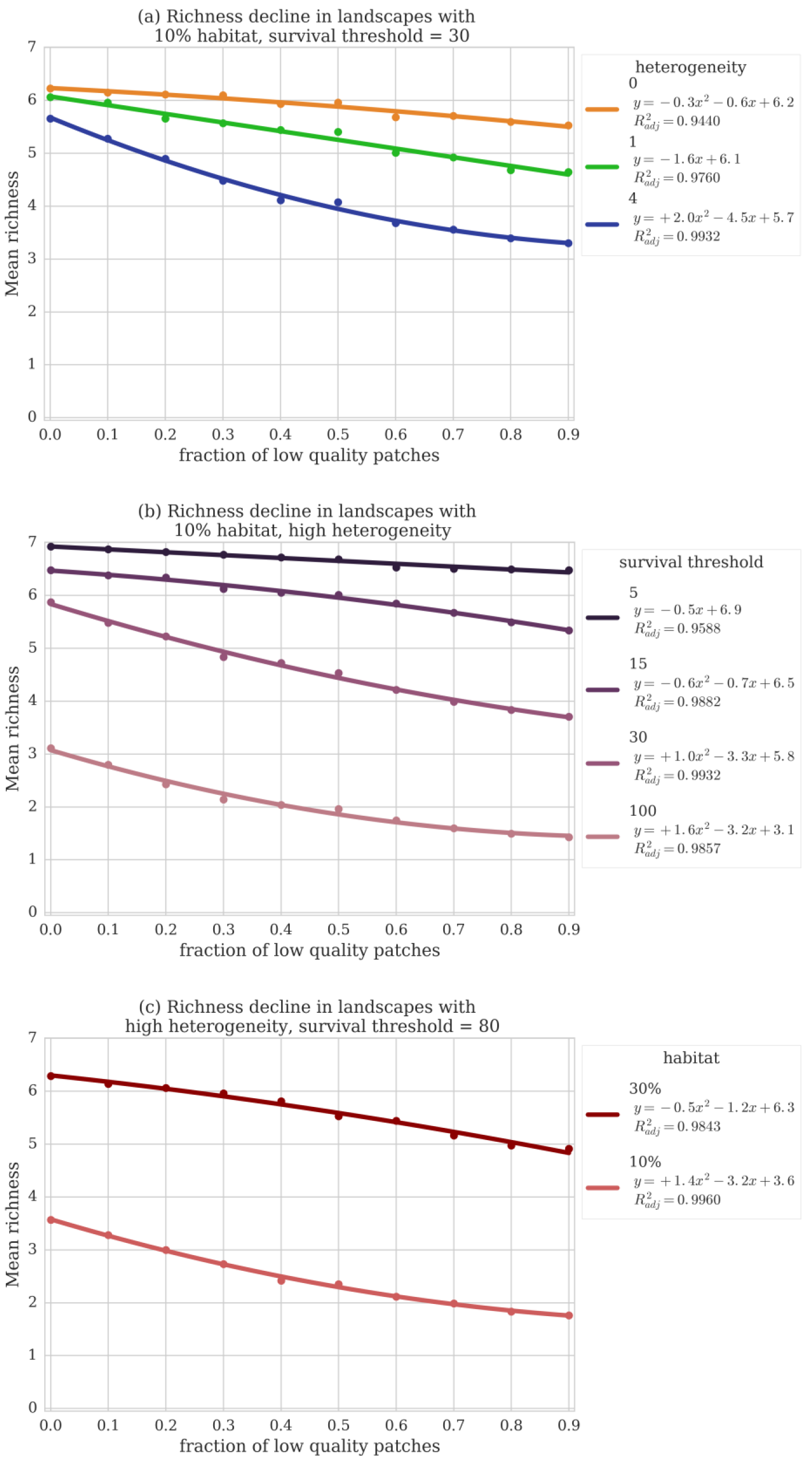
Factors that shape biodiversity decline along an intensification gradient. Richness decline along an intensification gradient shifts from a concave down to a concave up shape as stresses on biodiversity increase, either by (a) augmenting landscape heterogeneity, (b) applying stricter survival thresholds to species or (c) reducing habitat area.

In order to illustrate how the matrix facilitates or restricts a metacommunity dynamic through migration, we plotted the typical spatial distribution of richness in 10%-habitat landscapes reached by a single community in our experiments (using *survival threshold* = 0.1 for each patch), (Figure 5). At low intensification, richness is almost uniformly distributed across the landscapes, suggesting that habitat patches are coupled in a metacommunity dynamic. In contrast, at high intensification, habitat patches become isolated, though species survive inside them (see discussion on heterogeneity).

**Figure 5.**
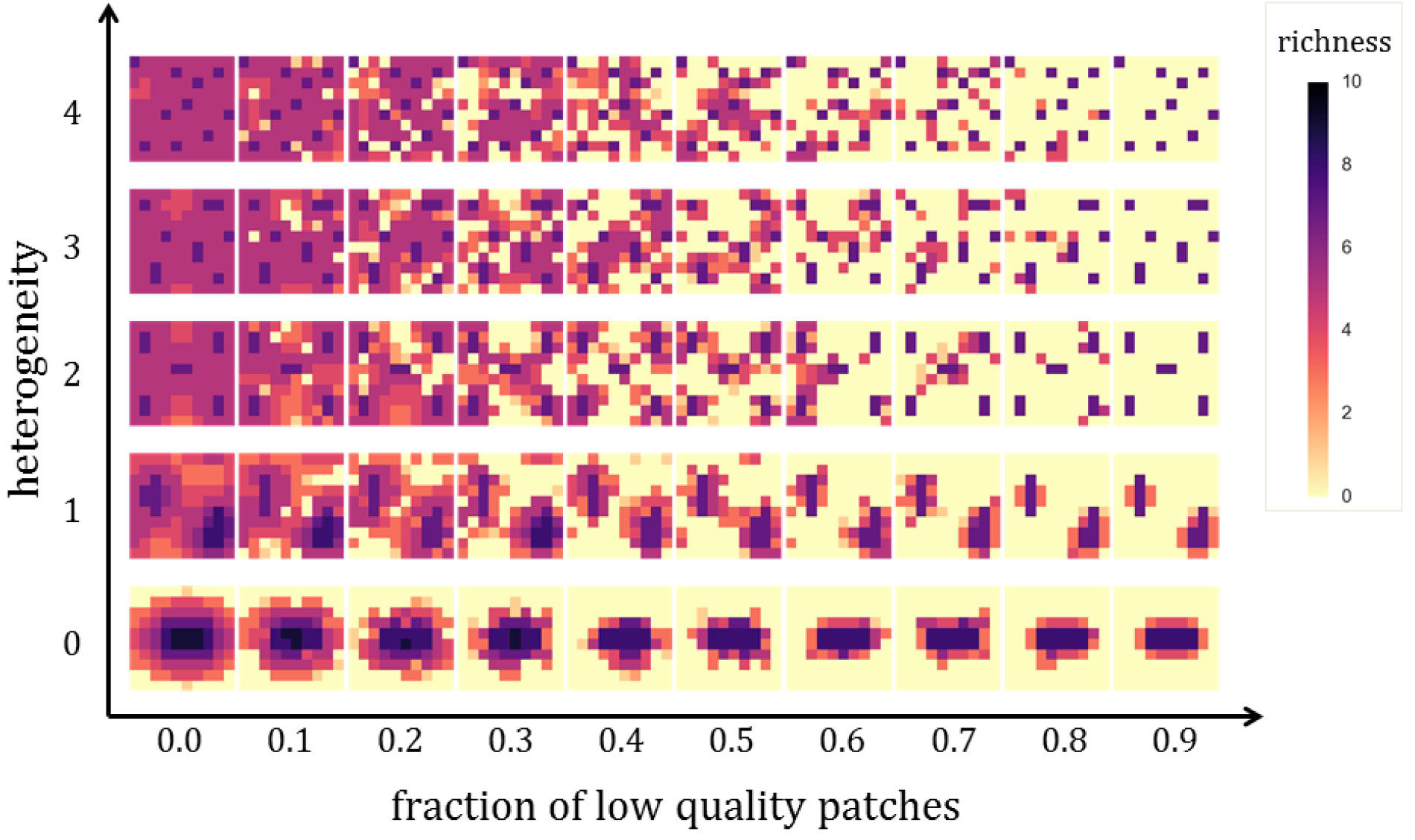
Spatial distribution of richness. Typical spatial distribution of richness in 10%-habitat landscapes reached by a single community in our experiments (*survival threshold* = 0.1 for each patch). The matrix facilitates or restricts a metacommunity dynamic: at low intensification, richness is almost uniformly distributed across the landscapes, suggesting that habitat patches are coupled in a metacommunity dynamic; in contrast, at high intensification, patches become isolated, thus increasing biodiversity’s vulnerability to potential perturbations. See Supplementary Figure 4 for scenario with 30% habitat.

### Effect of species’ survival thresholds on the shape of richness decline curves

We analyzed the data for high heterogeneity (level 2) varying the survival threshold for populations in the hypothetical communities by increasing orders of magnitude ranging from 0.0001 to 100, with a detailed analysis in the interval from 1 to 100 (Figure 4b). Below there is not a substantial loss in richness, and it generally follows a linear decline with a slight slope. Next, within the interval *survival threshold* = 10 to *survival threshold* = 25, richness curves gradually shift from a concave down to a concave up shape. Above *survival threshold* = 25, richness curves consistently show a concave up curve, but at the same time the proportion of communities that are extinct on initial conditions increases. These trends are similar for other heterogeneity levels, though the specific threshold values where the shifts occur differ.

### Effect of habitat proportion on the shape of richness decline curves

We repeated the previous analyses for a scenario where landscapes have 30% of habitat patches (Supplementary Figure 2). Here we use *survival threshold* = 80. In low heterogeneity levels (levels 0 and 1), richness declines linearly, though the change is small. Then, in heterogeneity levels 2 and 3, richness declines as a concave down curve; in heterogeneity level 4, richness declines linearly, but there is not a significant difference in richness distributions between levels 2, 3 and 4 at either degree of intensification. Moreover, there is not a significant difference between heterogeneity levels below 40% of intensification, and above 40% of intensification there are two statistically distinct richness distributions associated to high (levels 4, 3 and 2) and low (levels 1 and 0) heterogeneity levels, which contrasts with the three distinct responses found for 10%-habitat landscapes. While the effect of habitat proportion can be clearly observed considering a *survival threshold* = 80, the trends mentioned above do not depend on this particular value.

Comparing richness response at the same heterogeneity level highlights the effect of habitat area on the curve shape. We consider, for example, richness decline at heterogeneity level 2 in both 10% and 30% habitat areas (Figure 4c): in contrast with 10%-habitat landscapes, richness is notably higher in 30%-habitat landscapes and declines as a concave down curve as matrix quality decreases. Again, this suggests that a shift in the concavity of richness decline can occur as a result of reducing habitat area.

## DISCUSSION

In this work we analyzed factors that shape biodiversity’s response to agricultural intensification. We found that richness decline along an intensification gradient shifts from a concave down to a concave up curve, this is, from a more robust to a less robust biodiversity response, as landscape heterogeneity increases, as stricter survival thresholds are applied to species or as habitat area is reduced. Our work addresses some of the simplifications that make the land-sparing/land-sharing dichotomy a limited framework for comparing landscape management strategies. In particular, we studied species richness in response to the intensification of agricultural management, rather than yield, considering both the dynamics of ecological communities and landscapes explicitly defined in a combined gradient in matrix quality and configurational heterogeneity of habitat patches.

### Heterogeneity

Our results suggest that in landscapes with a small amount of habitat (10%), biodiversity is robust to agricultural intensification when habitat presents low heterogeneity (i.e. patches are clustered and contiguous instead of dispersed); in contrast, there is a significant impact of intensification on biodiversity in high heterogeneity landscapes, hence they also present less richness than low heterogeneity ones. The effect of heterogeneity on biodiversity becomes more important as intensification increases: biodiversity is similar in landscapes with highest matrix quality, irrespective of habitat configuration, but at higher degrees of intensification, there are distinct responses of biodiversity to high and low heterogeneity. In our model, this negative effect of heterogeneity on biodiversity is due to species spending a larger amount of time in the matrix, which increases overall mortality rate.

Chaplin-Kramer and collaborators (2015) report a similar shift in the concavity of biodiversity response to agricultural expansion induced by forest fragmentation in realistic landscapes. They found that while mean species abundance is robust when agricultural expansion advances from forest edge in (low fragmentation), it declines rapidly in a scenario of maximal forest fragmentation. Although the focus is on habitat loss rather than intensification, these distinct responses to spatial patterns of habitat are qualitatively consistent with our results.

While in this work we consider identical dispersal rates for all species, in a theoretical study that took into account both species’ dispersal abilities and landscapes with varying habitat quality comparable to the decrease in matrix quality in our model, the authors also found a negative effect of fragmentation on population size (Wiegand et al. 2005). Furthermore, in scenarios where species presented a metapopulation dynamic, fragmentation did not have an effect on population size in functionally connected landscapes (high quality matrix), but it had a large impact in landscapes with low quality matrix. This agrees with our results in that the effect of heterogeneity on biodiversity becomes more important as intensification increases.

Our results might overestimate the robustness of biodiversity’s response to intensification in low heterogeneity landscapes. We expect that in these landscapes the isolation of habitat patches at high degrees of intensification (Figure 5) would increase biodiversity’s vulnerability if we introduced a local extinction probability (for example, as a function of patch size, like it is assumed in classical metapopulation models); then, despite supporting greater richness, a potential perturbation could lead to local extinctions without the possibility of recolonization from other patches. In a similar way, Kremen (2015) suggested that empirical studies favoring land-sparing might also overestimate biodiversity persistence since their limited temporal scale ignores time lags for species extinctions that result from the isolation of habitat patches in low-quality matrices. Omitting local extinction probabilities allows us to focus on the effect of fragmentation and matrix quality on biodiversity decline, though it remains an important simplification in our model.

### Survival threshold

We found that increasing species’ survival threshold, which represents the minimum viable population in our model, gradually turns biodiversity response to intensification from a robust response to an abrupt decline. Because population persistence depends to some extent on population size (Shaffer 1981), it is reasonable to expect that imposing stricter conditions for species’ survival generates the qualitatively different responses of biodiversity to intensification we found. Our results suggest that the range of values for the survival threshold where biodiversity presents a robust response to intensification is smaller than that of an abrupt decline. Nevertheless, further analysis would require setting species-specific survival thresholds to reflect that the minimum viable population highly depends on species attributes and environmental conditions. See Larsen and Noack (2017) for an example where responses to agricultural practices depend more on individual attributes (of both crops and pests) than on landscape characteristics.

### Habitat

We found that reducing habitat area (from 30% to 10%) amplifies the impact of intensification on biodiversity. Landscapes with 10%-habitat area presented not only less richness than landscapes with the higher proportion of habitat, but a tendency towards abrupt biodiversity decline in response to intensification that contrasts with the robust response found for 30%-habitat landscapes (Figure 4c). In our model, the negative effect of habitat loss on biodiversity is mainly due to the obvious reduction in suitable area for species’ reproduction, which is a well-known cause of biodiversity decline (MacArthur & Wilson 2015). Another, indirect, effect of habitat loss, especially in high heterogeneity landscapes, is the increased distance between habitat patches that, similarly to fragmentation, affects the time spent in the matrix, and thus, overall mortality rate.

In agreement with our results, Batáry and collaborators (2011) found that the effect of landscape quality is stronger in scenarios with less natural habitat. In addition, the authors highlight the role of species’ dispersal rates in determining the effective distance between patches. Species with high dispersal rates are less affected by matrix quality, for instance, the high mobility of birds allows them to locate and exploit fields of high resource, independent of landscape features (Tscharntke et al. 2005, Batáry et al. 2011). On the other hand, species with low dispersal rates spend more time in between habitat patches, making them susceptible to changes in agricultural management (Thomas 2000). Therefore dispersal rates contribute to the effect of both habitat area and heterogeneity.

### Limitations

As mentioned above, considering equal survival thresholds and dispersal rates for all species is a limitation in our model. This, in turn, is a consequence of not assigning specific identities to hypothetical species. Choosing these parameters according to species’ traits would make populations differentially vulnerable to intensification and other perturbations, but we cannot extrapolate from our results how it would affect community and migration dynamics. Nonetheless, the lack of information for calibrating a model with hypothetical communities makes these assumptions the most parsimonious (González González et al. 2016). Besides, they allow us to emphasize the general effects of matrix quality and heterogeneity over those of species’ traits.

### Concluding remarks

At first sight, our results interpreted from the land-sparing/land-sharing framework would imply that land-sparing is preferable in high-heterogeneity landscapes with very little habitat whereas land-sharing is better in landscapes with greater habitat area. But this might be in conflict with the widespread association of land-sharing to high-heterogeneity landscapes and land-sparing to high-habitat ones. A conceptual gap in the land-sparing/land-sharing debate with respect to the characterization of landscape heterogeneity underlies this inconsistency and limits the comparison of management strategies. It has been argued that an ideal strategy would combine features of both sharing and sparing (Butsic and Kuemmerle 2015, Kremen 2015); however, locating management strategies within a continuum requires, in the first place, defining such continuum in the variables of interest.

We expect that our findings and the proposed model contribute to a nuanced understanding of how different landscape and ecological factors shape biodiversity response to agricultural intensification. In particular, our work illustrates the interdependent effects of heterogeneity, habitat availability and matrix quality. This could inform the design of management strategies that consider the impacts on livelihoods as much as on biodiversity. If a socially desirable strategy imposed constraints on landscape composition or configuration, then our theoretical approach could help explore complementary landscape components that could be managed to support a robust response of biodiversity. For example, beyond the land-sharing/land-sparing debate, it has been argued that small-scale agroecological strategies are key to achieve food sovereignty, which considers the preservation of biocultural heritage, high-quality soil and clean water availability, low-input food production and distribution, and the associated human rights to health, adequate and safe food, land and cultural identity (De Schutter 2011, Chappell et al. 2013, Holt-Giménez and Altieri 2013, Benett 2017, Gonzalez-Ortega et al. 2017). If such small-scale agroecological strategies were favored, one could ask what landscape elements could be managed to balance conservation and production goals under this scenario. Thus, rather than recommending general guidelines, we provide tools for the discussion of integrative and sustainable management strategies.

## ACKNOWLEDGMENTS

The authors thank Lev Jardón-Barbolla and members of *La Parcela* Laboratory for their valuable comments and suggestions. Cecilia González González and Ana L. Urrutia acknowledge the graduate program “Posgrado en Ciencias Biológicas, Universidad Nacional Autónoma de México”. This article covers part of the requirements to obtain the M.Sc. Degree in Biological Sciences (Ecology). Irene Ramos acknowledges the graduate program “Posgrado en Ciencias de la Sostenibilidad, Universidad Nacional Autónoma de México”. M.Benítez acknowledges financial support from UNAM-DGAPA-PAPIIT (IA200714; IN113013; IA202515) and CONACyT (221341; 247672).

## SUPPLEMENTARY INFORMATION

### Richness decline in 10%-habitat landscapes

**Figure S1.**
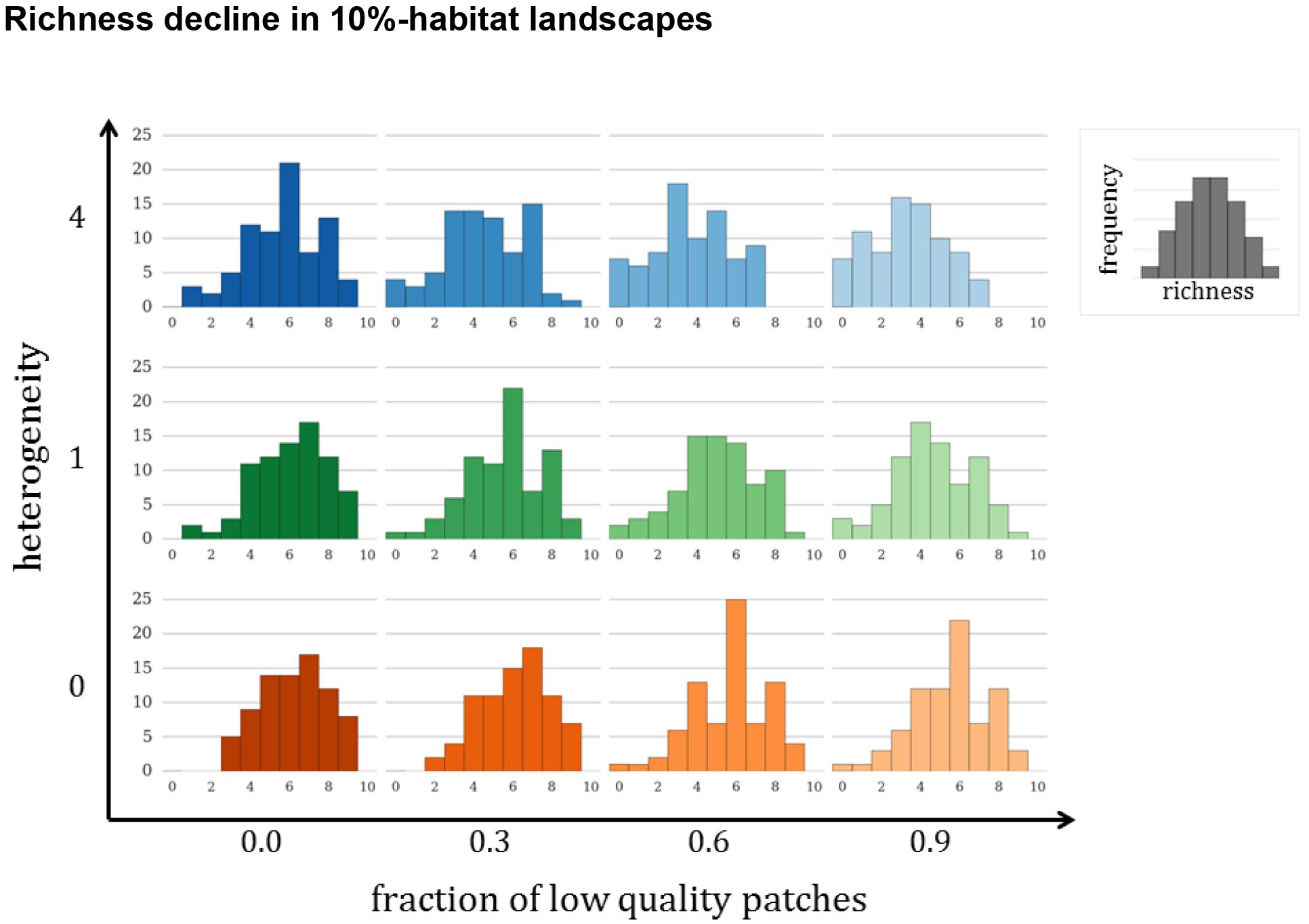
Frequency distributions of species richness that were reached by the 79 analyzed communities in 10%-habitat landscapes. At each intensification and heterogeneity combination, histograms show how many communities reached a given richness. We show selected points across the intensification gradient and the three heterogeneity levels that presented distinct responses (i.e. richness distributions showed significant differences).

### Richness decline in 30%-habitat landscapes

**Figure S2.**
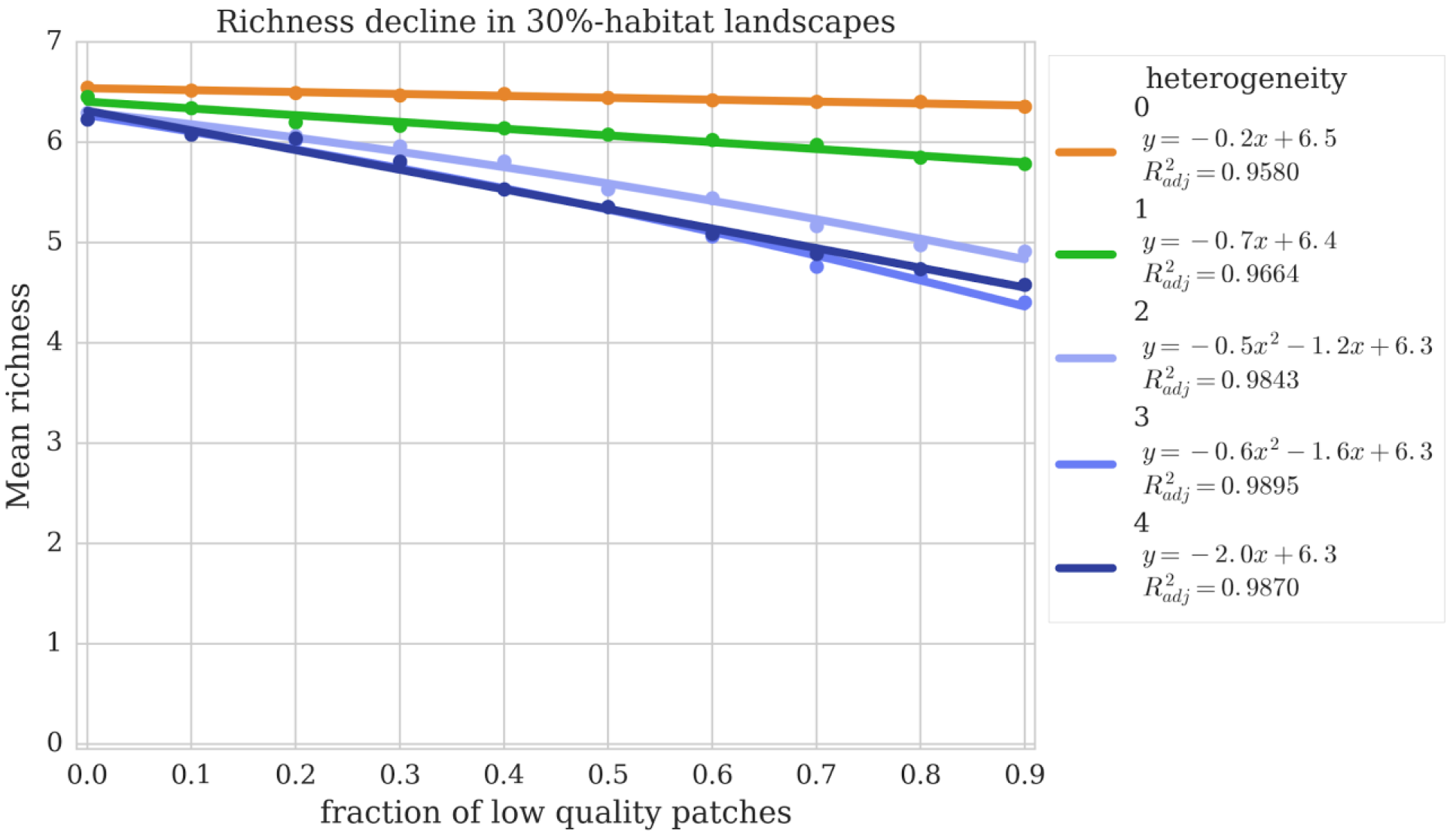
Richness decline along an intensification gradient (x-axis) in 30%-habitat landscapes (*survival threshold* = 80). In low heterogeneity landscapes (level 0 and 1), richness declines linearly, though the change is small. In contrast, richness follows a concave down curve in high heterogeneity landscapes (levels 2 and 3); in heterogeneity level 4, richness declines linearly, but there is not a significant difference in richness distributions between levels 2, 3 and 4 at either degree of intensification. Each data point is the average richness of 79 communities.

**Figure S3.**
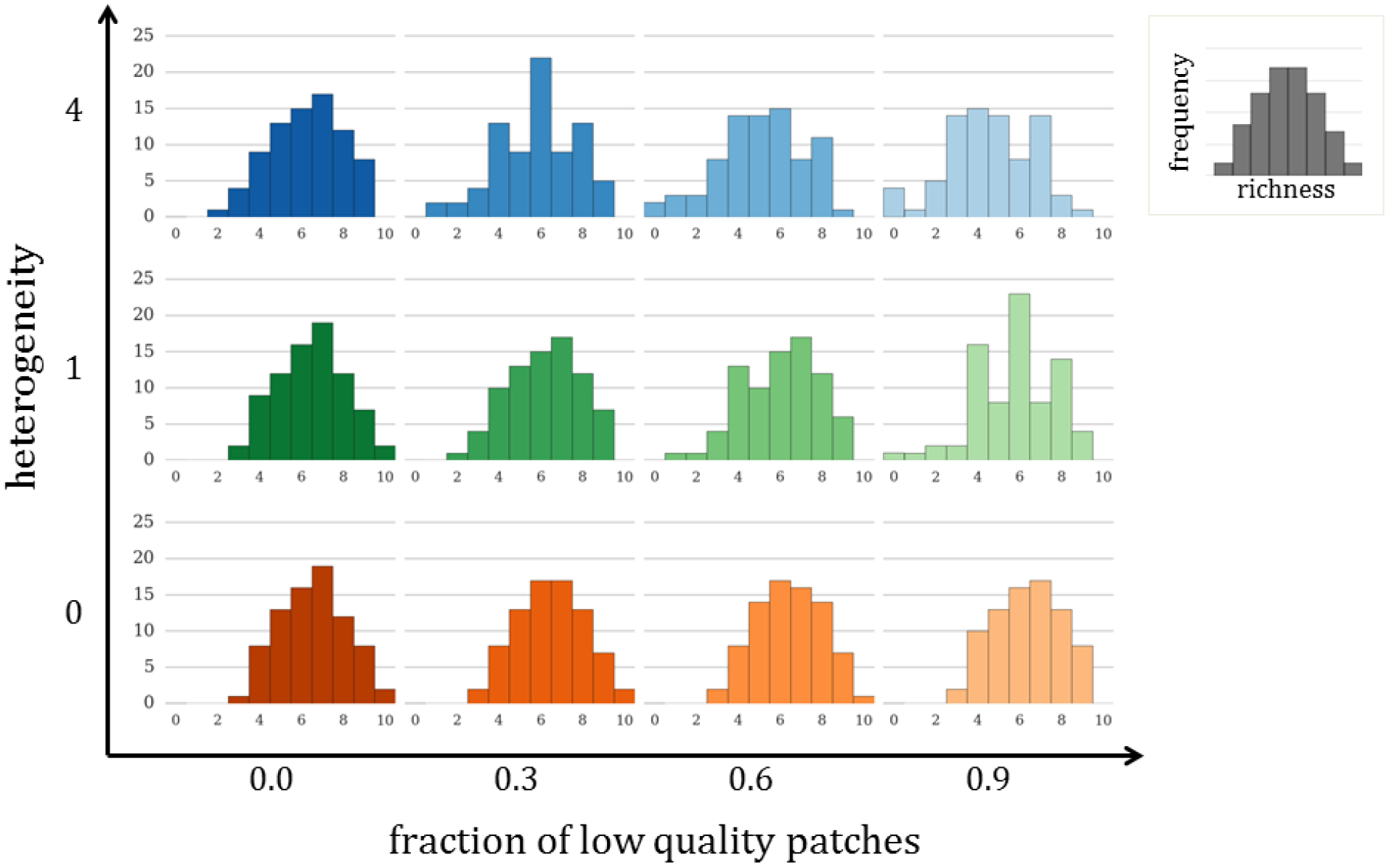
Frequency distributions of species richness that were reached by the 79 analyzed communities in 30%-habitat landscapes. Above 40% of intensification there are two statistically distinct richness distributions associated to high (levels 4, 3 and 2) and low (levels 1 and 0) heterogeneity levels, which contrasts with the three distinct responses found for 10%-habitat landscapes.

**Figure S4.**
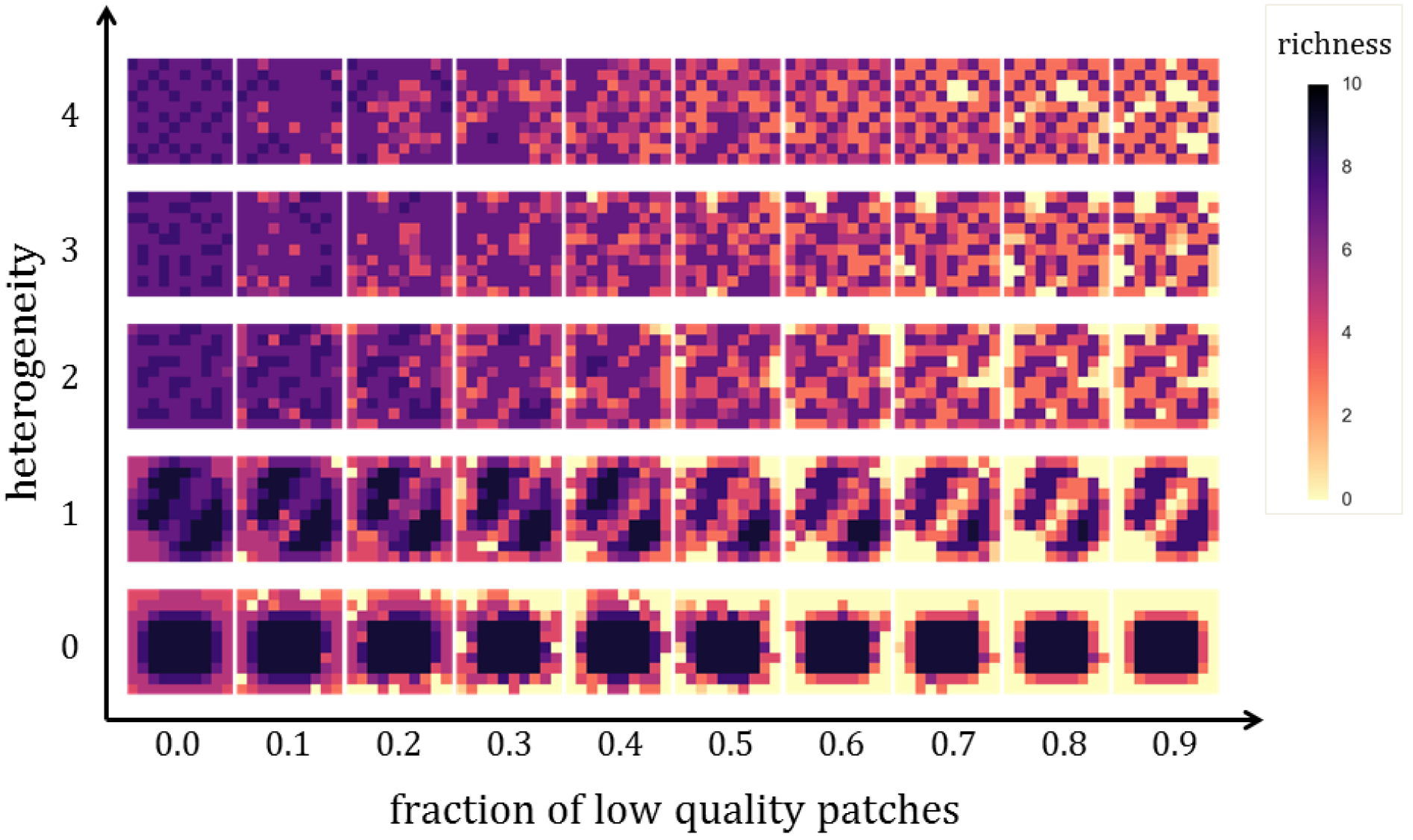
Typical spatial distribution of richness in 30%-habitat landscapes reached by a single community in our experiments (*survival threshold* = 0.1 for each patch). The effect of intensification on richness is not as severe as in 10%-habitat landscapes because there is more suitable habitat for reproduction and, in the case of high heterogeneity landscapes, because the distance between patches is shorter, thus species spend less time in the matrix.

